# *De novo* Prediction of Cell-Drug Sensitivities Using Deep Learning-based Graph Regularized Matrix Factorization

**DOI:** 10.1101/2021.10.11.462450

**Authors:** Shuangxia Ren, Yifeng Tao, Ke Yu, Yifan Xue, Russell Schwartz, Xinghua Lu

**Author notes:** Both authors contributed equally to this work (Shuangxia Ren, Yifeng Tao).

## Abstract

Application of artificial intelligence (AI) in precision oncology typically involves predicting whether the cancer cells of a patient (previously unseen by AI models) will respond to any of a set of existing anticancer drugs, based on responses of previous training cell samples to those drugs. To expand the repertoire of anticancer drugs, AI has also been used to repurpose drugs that have not been tested in an anticancer setting, i.e., predicting the anticancer effects of a new drug on previously unseen cancer cells *de novo*. Here, we report a computational model that addresses both of the above tasks in a unified AI framework. Our model, referred to as deep learning-based graph regularized matrix factorization (DeepGRMF), integrates neural networks, graph models, and matrix-factorization techniques to utilize diverse information from drug chemical structures, their impact on cellular signaling systems, and cancer cell cellular states to predict cell response to drugs. DeepGRMF learns embeddings of drugs so that drugs sharing similar structures and mechanisms of action (MOAs) are closely related in the embedding space. Similarly, DeepGRMF also learns representation embeddings of cells such that cells sharing similar cellular states and drug responses are closely related. Evaluation of DeepGRMF and competing models on Genomics of Drug Sensitivity in Cancer (GDSC) and Cancer Cell Line Encyclopedia (CCLE) datasets show its superiority in prediction performance. Finally, we show that the model is capable of predicting effectiveness of a chemotherapy regimen on patient outcomes for the lung cancer patients in The Cancer Genome Atlas (TCGA) dataset.^*^

## 1. Introduction

Precision oncology aims to treat each patient with an individually tailored therapy regimen to achieve better outcomes and minimize side effects.^1^ Currently, a common practice in precision oncology is to prescribe molecularly targeted drugs that are intended specifically to counteract aberrant signals resulting from tumor-specific genomic alterations. However, such genome-driven precision oncology has so far covered a limited fraction of patients.^2^ Furthermore, responses to specific targeted therapeutics are often short-lived due to tumor heterogeneity and development of resistance.^3,4^ Currently, most cancer patients are treated with “standard chemotherapies” that are not personalized, and a large proportion of patients do not respond to these therapies but suffer the full brunt of their side effects.

Therefore, the success of precision oncology requires the capability to accurately predict the responses of a patient’s cancer cells to existing anti-cancer drugs and select an optimal regimen, potentially adjusting over time in response to incipient resistance. The resulting need for an expanded repertoire of anticancer drugs has led to an active research effort to discover and repurpose FDA-approved drugs that are not yet considered as anticancer therapeutics but may function as such. Furthermore, anticancer therapies often involve a combination of multiple drugs, and the large number of possible combinations of anticancer drugs prevents systematic clinical trials to develop novel therapies. The above unmet needs call for methods for predicting effects of drugs on cancer cells even when they have not been tested in such a setting, i.e., *de novo* prediction of drug effects. Contemporary large-scale pharmacogenomic studies, such as Genomics of Drug Sensitivity in Cancer (GDSC),^5^ Cancer Cell Line Encyclopedia (CCLE),^6^ The Cancer Genome Atlas (TCGA),^7^ Library of Integrated Network-based Cellular Signatures (LINCS)^8,9^ provide valuable information for exploring the above directions, but would benefit greatly from computational systems capable of mining the information they provide and using it to make accurate prediction about potential new therapeutic regimens and thus advance precision oncology.

Prediction of cell-drug responses can be formulated as a recommendation problem (e.g., collaborative filtering^10^). More specifically, given information regarding a collection of cancer cells (e.g., genomic and transcriptomic profiles of different cancer cell lines) and their responses to different drugs, we would like to learn the representations of the cells such that cells sharing similar representations respond similarly to drugs. Similarly, given information regarding a collection of drugs (e.g., chemical structures and knowledge regarding the drugs) and their effects on different cancer cells, we would like to learn representations of the drugs such that drugs sharing similar representations have similar effects on the cells. After training, when provided with information of a new sample of cells, a recommendation system should be able to predict response of each cancer cell to different drugs. Alternatively, provided information on a new drug, the system should be able to predict the effects of the drug on different cells. Finally, given a new cell sample and a new drug (both previously unseen in training process), the recommendation system should be able to map the cell and/or the drug to respective representations and predict the cell-drug response *de novo*.

A variety of computational methods have been developed to predict the drug sensitivities of cancer cell lines to a large number of drugs.^11–13^ However, the majority of previous drug-sensitivity models concentrate on predicting responses of different cells to an individual drug, and few have attempted to address the problem as posed above. These models do not fully take advantage of available information on other drugs with respect to cells to learn from drugs with similar chemical structures or mechanisms of action (MOAs), nor do they take advantage of the fact that some cancer cells share similar drug response profiles to learn common representations of such cells. Prior approaches to learning representations of drugs have transformed their chemical structures from a form defined by the simplified molecular-input line-entry system (SMILES) into a vector (an embedding) that can be concatenated with cell embeddings in a deep learning model to predict drug response.^14–16^ However, this approach does not utilize a rich body of information regarding the functional impact of chemicals on cell signaling systems,^8,9,17^ which is highly relevant to the MOAs of drugs^18^ and thus relevant to predicting drug responses.

In this study, we developed a method called DeepGRMF (deep learning-based graph regularized matrix factorization). The main innovation of our model lies in integration of multiple sources of information and different learning techniques, including: 1) **Integrative representation of drugs**. For the representation of drugs, we combined three kinds of information regarding a drug — its chemical structure,^19^ its impact on cellular transcriptomic signaling,^17^ and its pathway information^20^ — to make the representation more informative. 2) **Representation learning through collaborative filtering**. Our model employs a framework of collaborative filtering based on matrix-factorization, which learns representations of cells based on the shared responses with respect to drugs as well as representations of drugs based on their common effects on cells. 3) **Graph-based regularization**. We adopted a graph-based regularization approach^21^ to enhance the performance of collaborative filter modeling of cell-drug responses. 4) **Neural-network-based mapping from raw data to cell and drug factor matrix**. We incorporated two neural network models which map cells (or drugs) to their corresponding factor matrix. This enables us to perform *de novo* prediction of responses between a pair of previously unseen cell and drug.

Our results indicate that each of the above steps individually enhanced overall performance, and that the complete model outperforms current state-of-the-art models in predicting drug responses.

## 2. Materials and methods

### 2.1. Data pre-processing

We collected the pharmacogenomic data from GDSC (https://www.cancerrxgene.org) and CCLE (https://portals.broadinstitute.org/ccle) to train and test models for predicting drug cell-drug responses. We used the method described in previous work^11^ to process the gene expression data and drug sensitivity data. In brief, the gene expression data were normalized by the robust multi-array averaging, and genes with high variances were identified by medium variance analysis, bimodal mixture fitting, and statistical significance of modes. After filtering, we applied a nonparanormal transformation for distribution normalization and a min-max normalization to normalize the value of expression data in a range between 0 to 1. Finally, we retained GDSC and CCLE datasets containing gene expression data of 2,758 genes in 954 and 477 cell lines separately. For drug sensitivity data, activity area (AA) was used. To facilitate application in clinical practice, we discretized the continuous value into two categories, sensitive (one) and resistant (zero).^11^ Since drug embedding is based on SMILES strings, we only selected drugs with known SMILES strings, which resulted in 301 drugs in GDSC and 24 drugs in CCLE (16 drugs existing in GDSC and 8 new drugs). For the drug pathway information, the GDSC dataset has already provided a type of pathway name for each drug. For the new drug in CCLE, we labeled its pathway name manually using the pathway name in GDSC.

For predicting effectiveness of a chemotherapy regimen on real patients using the cell-line trained drug sensitivity prediction model, we collected RNAseq expression data of lung adenocarcinoma (LUAD) and lung squamous cell carcinoma (LUSC) patients from the UCSC Xena TCGA data portal (https://xena.ucsc.edu). The gene expression data in TCGA was processed using the same procedure of processing the gene expression data in GDSC and CCLE. The drug usage data of 179 LUAD and 144 lung squamous cell carcinoma LUSC patients were downloaded from the Genomic Data Commons Data Portal (https://portal.gdc.cancer.gov). We downloaded the corresponding survival data from the UCSC Xena data portal. We kept data of patients who received adjuvant therapies. This resulted in a lung cancer test dataset with drug usage and survival information for 182 patients. Since we focused on four drugs, cisplatin, pemetrexed, paclitaxel, and vinorelbine, finally we had a dataset with 62 adjuvant LUAD and LUSC patients.

### 2.2. Learning integrative drug embedding

To obtain an integrative drug embedding (IDE) reflecting multiple aspects of the drug, we adopted a semi-supervised method^19^ that integrated two sources of information: 1) chemical structure of the drug, and 2) functional impact of the drug on gene expression.^17^ To represent chemical structures, we used the SMILES^22^ representation of molecules and obtained SMILES strings of 250K drug-like molecules from the ZINC^23^ database. To represent the functional impact on gene expression, we trained a variational auto-encoder (VAE)^24^ model on the transcriptomic data of cell lines treated by different drugs from the LINCS database.^8^ Based on the assumption that the cellular transcriptomic profile in response to a drug treatment reflects the MOAs of the drug on the cell, we learned the MOAs representations of 1,825 drugs.^17^ After obtaining the two sources of information, we used a VAE to encode the SMILES strings of molecules^19^ and utilized the drug-drug similarity computed from their MOAs representations to regularize the drug embedding space so that drugs with similar MOAs profiles were clustered together. Note that our approach allows us to obtain the IDE of a new drug by mapping its SMILES string to the pre-trained drug embedding space.

### 2.3. Model architecture

The overall architecture of DeepGRMF is shown in Fig. 1. This model has two modules: 1) a graph regularized matrix factorization that decomposes the drug response matrix into the product of two lower dimensional matrices, i.e., cell line factor matrix *A* and drug factor matrix *B*; 2) two neural networks that learn the functions that map the representations of cell line and drug to their hidden factors, i.e., *C* → *A* and *D* ∪ *E* → *B*, respectively.

**Fig. 1:**
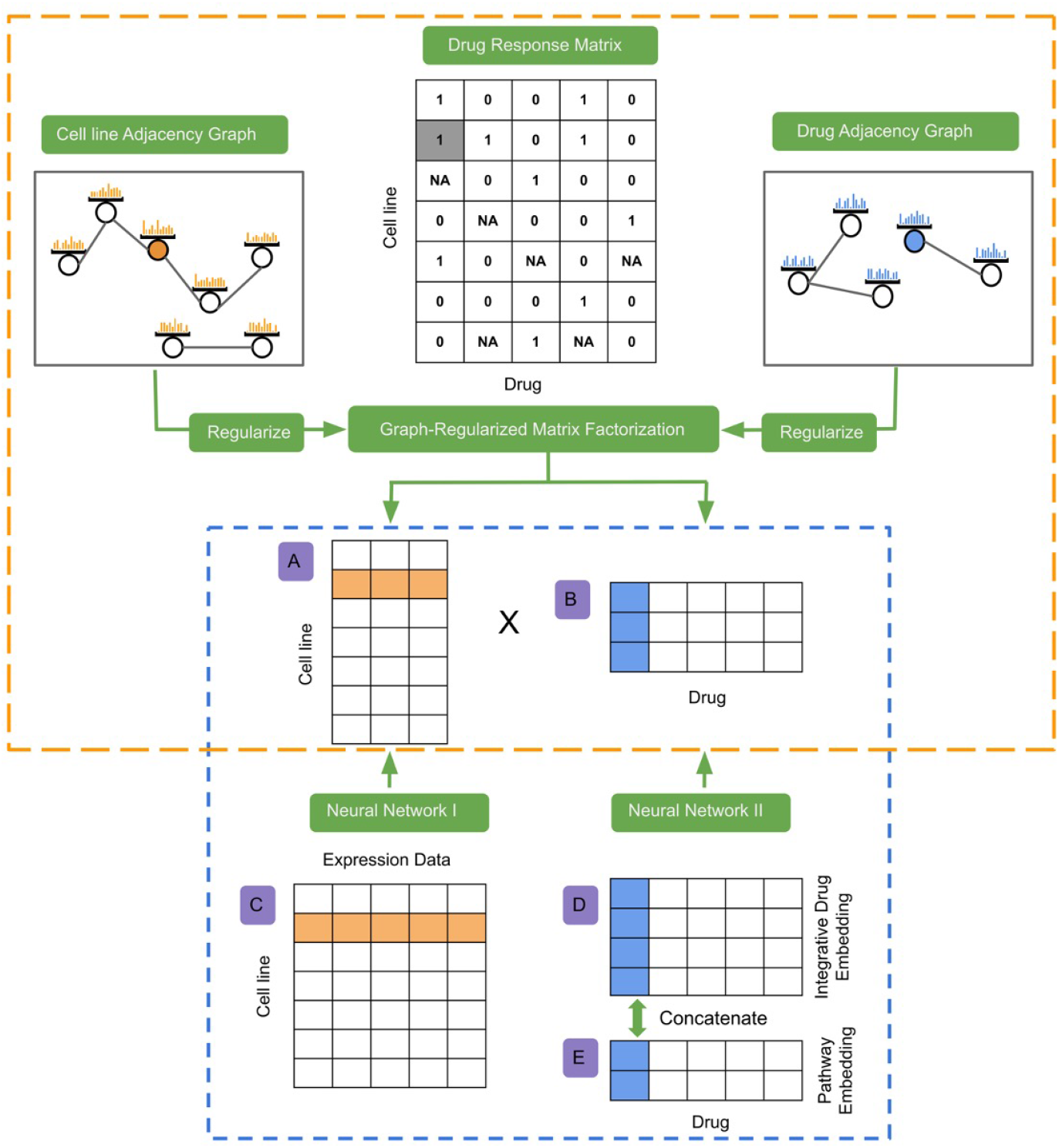
Diagram of DeepGRMF model. 1) The orange dotted box shows the procedure of the first module: using a graph regularized matrix factorization to decompose the drug response matrix into the product of cell line factor matrix *A* and drug factor matrix *B*. 2) The blue dotted box shows the procedure of the second module: using two separate neural networks to learn the mapping functions. The neural network I is used to learn the mapping function for cell lines, which maps gene expression matrix *C* to cell line factor matrix *A*. The neural network II is used to learn the mapping function for drugs, which maps the drug embedding obtained by concatenating integrative drug embedding matrix *D* and pathway embedding matrix *E* to drug factor matrix *B*.

#### 2.3.1. Graph-regularized matrix factorization

We used matrix factorization to decompose the drug response matrix, *Y* ∈ ℝ^*n*×*m*^ into the cell line factor matrix, *A* ∈ ℝ^*n*×*d*^ and the drug factor matrix, *B* ∈ ℝ^*d*×*m*^. The variables *n*, *m* and *d* indicate the number of cell lines, the number of drugs, and the latent dimension respectively. The representations of cells and drugs in factor matrices capture the similarity of cell responses to drugs and effects of drugs on cells, so that if two cell lines have similar representations, they would respond similarly to certain drugs, and vice versa. However, this encoding does not incorporate intrinsic information of cells (e.g., cellular states) or drugs (e.g., chemical structures). Inspired by the work of Guan et al,^21^ we employed graph-regularized terms so that the similarities among the latent vectors in *A* and *B* are consistent with the pairwise similarities derived from the cell line gene expression matrix *C* and the drug embedding by concatenating IDE matrix *D* and pathway embedding matrix *E* denoted as *D* ∪ *E*.

The graph-regularized matrix factorization can be formulated as an optimization problem with loss functions and constraints. For the constraints, we used both graph regularizations of cell lines and drugs. We created adjacency matrices for cell lines (*W*_cell_ ∈ ℝ^*n*×*n*^) and drugs (*W*_drug_ ∈ ℝ^*m*×*m*^), respectively. The adjacency matrix is a representative description of a graph structure in matrix form and the elements of it represent whether pairs of vertices are adjacent in the graph or not. In our experiment, *W*_cell_ was constructed from the gene expression matrix *C* to measure the affinity between cell lines. To create *W*_cell_, a kernel function^25^ (*S*_cell_)_*i,j*_ = exp(−*T*_*i,j*_/*σ*) was first applied to convert the Euclidean distance between a pair of gene expression profiles into a similarity score within the range of [0, 1], where *T*_*i,j*_ = ∥**x**_*i*_ – **x**_*j*_∥ is the Euclidean distance between expression profiles **x**_*i*_ and **x**_*j*_, and *σ* is the mean of all the elements in *T*. We used *S*_cell_ to identify the set of top *p*-nearest neighbors for each cell line, and we set entries of these neighbors in the adjacency matrix to 1, and rest to 0.

To derive the drug adjacency matrix (*W*_drug_), we first created a similarity matrix *S*_drug_ based on the IDE using similar procedures describing above to obtain *S*_cell_, then we added a value of 0.5 to the similarity score in *S*_drug_ if a pair of drugs are in a common pathway. We then created an adjacency matrix (*W*_drug_) by only keeping edges connecting the *p*-nearest neighbors of each drug.

Given a cell line factor 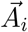 (*i*^th^ row of matrix *A*), and a drug factor 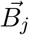 (*j*^th^ column of matrix *B*), we optimized the following loss function:

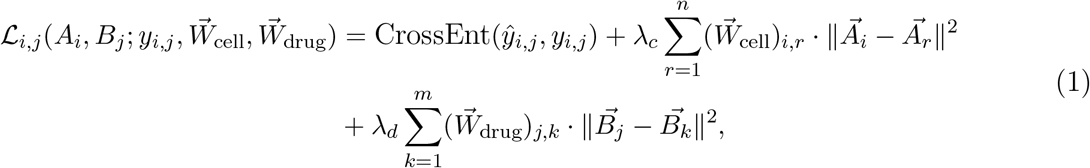

where 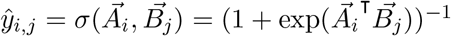 is the predicted sensitivity, *y_i,j_* is the ground truth sensitivity, and *λ_c_* and *λ_d_* are positive regularization weights. The loss function contains three terms. The first term is a cross-entropy loss. The second term is the graph regularization of cell lines, which enforces cell lines with similar gene expression to be close in the cell line factor space *A*. The last term is the graph regularization of drugs, which enforces drugs that are connected in the adjacency graph to be close in the d g factor space *B*. The final loss function is the sum of individual cell lines and drugs: 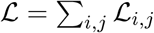.

At the training time, we implemented factor matrices A and B as two embedding layers with randomly initialized weights. Then we trained the graph-regularized matrix factorization module using Adaptive Moment Estimation (Adam) as gradient descent optimization algorithm (see 5.1 for implementation details). After the optimization was converged, we obtained the learned cell line factor matrix *A* and the learned drug factor matrix *B* and kept them as fixed during the training of the second module.

#### 2.3.2. Using neural networks to learn mapping function

To enable *de novo* prediction of responses between a pair of previously unseen cell and drug, we used neural networks to learn two mapping functions. Specifically, we used the neural network I (denoted as *f*_*θ*_) to map a cell line gene expression profiling 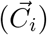 into its corresponding cell line factor 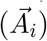. We used the neural network II (denoted as *f*_*ϕ*_) to map a drug’s embedding, which is concatenated by its IDE 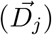 and pathway embedding 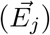, into its corresponding drug factor 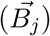. We adopted an embedding layer (denoted as *f*_*γ*_) to convert a drug’s pathway information 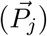 into its pathway embedding 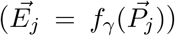. We trained these two neural networks separately and used the two loss functions for each network:

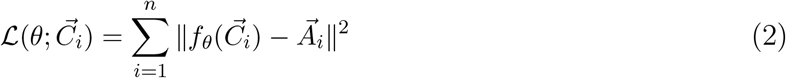

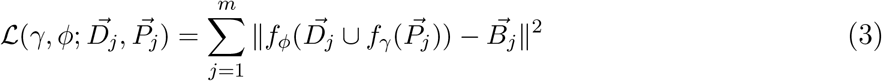

### 2.4. Evaluating drug sensitivity prediction on cell lines

We considered 3 scenarios of using our model to predict cell-drug responses: 1) Given a new cell line that has not been treated by any drug (Fig. A1**a**), we would apply our neural network I to predict its cell factor based on its transcriptomic profile and then apply collaborative filtering to predict its response to all the drugs. 2) Given a new drug that has not been tested on any cell line before, we would apply the neural network II to predict its drug factor based on its IDE and pathway information, and we would then apply collaborative filtering to predict its effects on all cells (Fig. A1**b**). Finally, 3) Given a new cell line and a new drug, we would first use neural network I and II to predict cell and drug factors respectively, and then apply collaborative filtering to predict the cell-drug response (Fig. A1**c**).

We used three schemes to evaluate the performance: 1) the disentangled performance of individual cell lines to all drugs (per-cell-line performance); 2) the disentangled performance of individual drugs to all cell lines (per-drug performance); 3) and the global performance ignoring the distinctions among cell lines and drugs (micro performance). We used area under the receiver operating characteristic (AUROC) and area under the precision-recall curve (AUPR) as evaluation metrics. We reported average per-cell-line and average per-drug AU-ROCs/AUPRs for comparisons.

### 2.5. Survival analysis on real patients

We evaluated performance of our model on real-world patients by first assigning patients into predicted *responders* and *non-responders*, and we then compared their survivals as a surrogate indicator of drug efficacy. From TCGA consortium, we collected clinical data, including treatments and overall survival, of 62 LUAD and LUSC patients, who received different combinations of cisplatin (41 cases), pemetrexed (19 cases), paclitaxel (17 cases), and vinorelbine (10 cases) as adjuvant therapies. We applied our model, which has been trained on GDSC dataset, to each patient to derive the probabilities of being sensitive to the drugs in the prescribed regimen. Since the probabilities for different drugs were not well calibrated, we assigned a patient as sensitive to a drug if the prediction probability for the patient is among the top 40^th^ percentile of all patients treated with the drug. We then designated a patient as a *responder* to a regimen if the patient is predicted to be sensitive to any of drugs in the regimen, otherwise as a *non-responder*. We tested our model on these drugs in two schemes: 1) Predicting efficacy of existing drugs to treat new cancer cells (previously unseen by models) as in Fig. A1**a**; 2) and predicting efficacy of new drugs (unseen during training) on new cancer cells as shown in Fig. A1**c**. For the second scheme, we removed these four drugs from GDSC dataset during training.

## 3. Results

### 3.1. Drug embedding analysis

We evaluated the quality of IDE and the contribution of each of its components, i.e., chemical structure, MOAs, and pathway information. Our evaluation was based on the heuristics that a pair of drugs with similar effects on cancer cells should be close in the drug embedding space. To that end, we calculated the drug-drug similarity (Jaccard coefficient) in terms of their cell-drug response profiles, and we computed the pairwise Euclidean distances between drug embeddings. For easy visualization, we divided drug-drug pairs equally into low (bottom 33%), medium, and high (top 33%) quantiles with respect to their Euclidean distances. Fig. 2A shows the relationship between these two pairwise measurements. We observed a decreasing trend (the blue curve) of drug sensitivity similarity from the low quantiles of Euclidean distance to the high quantiles of Euclidean distance in the drug embedding space using only chemical structure information. This trend (the orange curve) becomes more evident by using the MOAs as a regularization, suggesting that the drug embedding is augmented by adding the MOAs information. Similarly, Fig. 2B shows that including the pathway information into the IDE further boosted its quality.

**Fig. 2:**
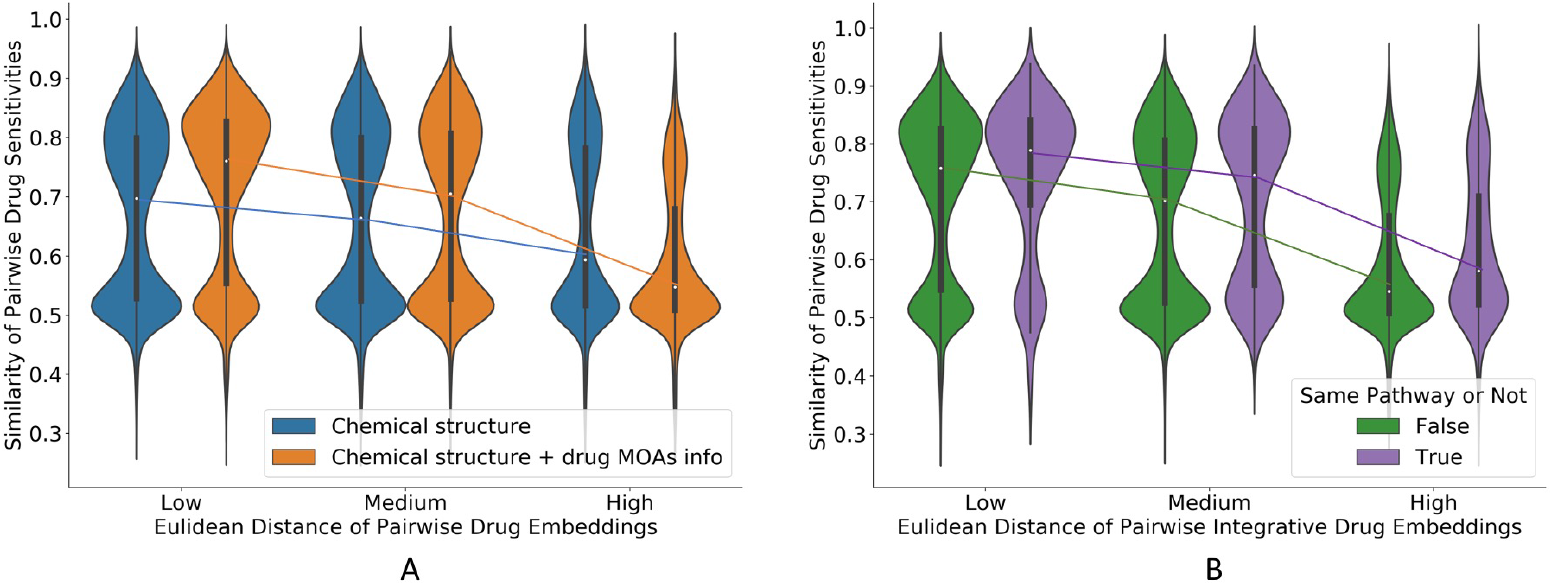
A) The relationship between similarities of drug sensitivity and Euclidean distance of drug representations using chemical structure with/without the drug MOAs information. B) The relationship between similarities of drug sensitivity and Euclidean distance of drug representations using chemical structure, drug effect with/without pathway information.

### 3.2. Drug sensitivity prediction on cell line

To evaluate the performance of DeepGRMF, we applied 25-fold cross-validations to GDSC dataset. As shown in Fig. A2, we adopted different train-test split strategies. In order to test the *out-of-sample* performance across different platforms or pipelines and examine the robustness of DeepGRMF, we also predicted and evaluated the drug response of unseen CCLE dataset with the *GDSC-trained* models.

#### 3.2.1. Drug sensitivity prediction of new cell lines to existing drugs

To predict drug sensitivity of new cell lines to existing drugs (Fig. A2**a**), the cell lines were split into 25 folds, every time we trained on 24 folds and tested on the remaining one. We evaluated the performance of DeepGRMF and compared it with two models: Lasso and DeepDSC,^15^ where Lasso is a classic model and DeepDSC is a state-of-the-art model to predict drug sensitivity. DeepGRMF outperformed these two models (all three were trained on GDSC data) in both GDSC and CCLE datasets (Table 1), indicating both better accuracy and generalization of the model. These results show the superiority of non-linear modeling for drug and gene expression in DeepGRMF over the linear modeling in Lasso. DeepDSC introduces non-linearity by concatenating the drug chemical features with cell line genomic features followed by a neural network to predict the drug sensitivity data. We assumed the collaborative filtering in DeepGRMF could better capture the interaction between cell line and drug.

**Table 1:** Performance of different models to predict drug response of new cell lines to existing drugs.

| Train/Val Data | Test Data | Model | Per Cell Line |  | Per Drug |  | Micro |  |
| --- | --- | --- | --- | --- | --- | --- | --- | --- |
|  |  |  | AUROC | AUPR | AUROC | AUPR | AUROC | AUPR |
| GDSC | GDSC | Lasso | 79.1 | 53.8 | 67.1 | 38.2 | 79.3 | 55.4 |
|  |  | DeepDSC | 80.0 | 54.8 | 67.7 | 38.8 | 79.9 | 56.4 |
|  |  | DeepGRMF | **83.2** | **60.1** | **70.9** | **41.8** | **83.1** | **62.0** |
| GDSC | CCLE | Lasso | 79.2 | 67.5 | 66.2 | 38.2 | 74.1 | 50.5 |
|  |  | DeepDSC | 80.0 | 68.3 | 67.0 | 40.5 | 75.1 | 51.5 |
|  |  | DeepGRMF | **82.0** | **70.9** | **67.9** | **41.6** | **76.0** | **53.7** |

#### 3.2.2. Drug sensitivity prediction of existing cell lines to new drugs

To predict drug sensitivity of existing cell lines to new drugs (Fig. A2**b**), we split drugs into 25 folds and used 24 folds for training and the remaining one for testing. The performance of DeepGRMF was compared to DeepDSC in the task of predicting drug sensitivity of existing cell lines to new drugs. DeepGRMF compared favorably to DeepDSC on both AUROC and AUPR (Table 2). We did not compare with Lasso because Lasso needs to build a different model for each cell line, while 1k cell lines in the dataset are too many. Since CCLE has different cell lines from GDSC, for this task we can not train on GDSC and test on CCLE.

**Table 2:** Performance of different models to predict drug response of existing cell lines to new drugs.

| Train/Val Data | Test Data | Model | Per Cell Line |  | Per Drug |  | Micro |  |
| --- | --- | --- | --- | --- | --- | --- | --- | --- |
|  |  |  | AUROC | AUPR | AUROC | AUPR | AUROC | AUPR |
| GDSC | GDSC | DeepDSC | 58.6 | 33.0 | 64.5 | 35.3 | 65.4 | 37.7 |
|  |  | DeepGRMF | **65.5** | **38.1** | **70.7** | **41.8** | **72.9** | **46.7** |

#### 3.2.3. Drug sensitivity prediction of new cell lines to new drugs

To predict drug sensitivity of new cell lines to new drugs (Fig. A2**c**), we used both new cell lines and new drugs to test the prediction performance. The cell lines and drugs were firstly split into *five* folds, and we then paired each drug fold with each cell line fold to create 25 folds in total. Every time we utilized one fold of cell line paired with one fold of drugs to test, the remaining pairs of cell lines and drugs were used to train the model.

Table 3 shows the comparison between DeepGRMF and DeepDSC in this evaluation scheme. DeepGRMF has better performance than DeepDSC on both AUROC and AUPR. Compared with DeepDSC, the graph regularization technique could further capture the similarity among cell lines and drugs, thus improving our performance.

**Table 3:** Performance of different models to predict drug response of new cell lines to new drugs.

| Train/Val Data | Test Data | Model | Per Cell Line |  | Per Drug |  | Micro |  |
| --- | --- | --- | --- | --- | --- | --- | --- | --- |
|  |  |  | AUROC | AUPR | AUROC | AUPR | AUROC | AUPR |
| GDSC | GDSC | DeepDSC | 58.2 | 31.6 | 55.6 | 28.5 | 59.8 | 31.9 |
|  |  | DeepGRMF | **64.6** | **36.6** | **61.4** | **33.8** | **66.9** | **38.9** |
| GDSC | CCLE | DeepDSC | 49.1 | 49.4 | 58.5 | 38.1 | 55.1 | 32.2 |
|  |  | DeepGRMF | **56.1** | **55.5** | **69.1** | **49.4** | **61.0** | **44.6** |

### 3.3. Survival analysis on real patients

As shown in Fig. 3, patients in the *responders* group survived significantly longer than the *non-responders* group regardless of whether we treated the four drugs (cisplatin, pemetrexed, paclitaxel, and vinorelbine) as existing drugs or new drugs. The log-rank test p-value is 0.03 and 0.05 for existing and new drugs, respectively. Our model successfully discriminated lung cancer patients into different drug response groups that are correlated with survival outcomes in both schemes. Compared with treating the four drugs as new drugs, treating the four drugs as existing drugs better separated the two groups.

**Fig. 3:**
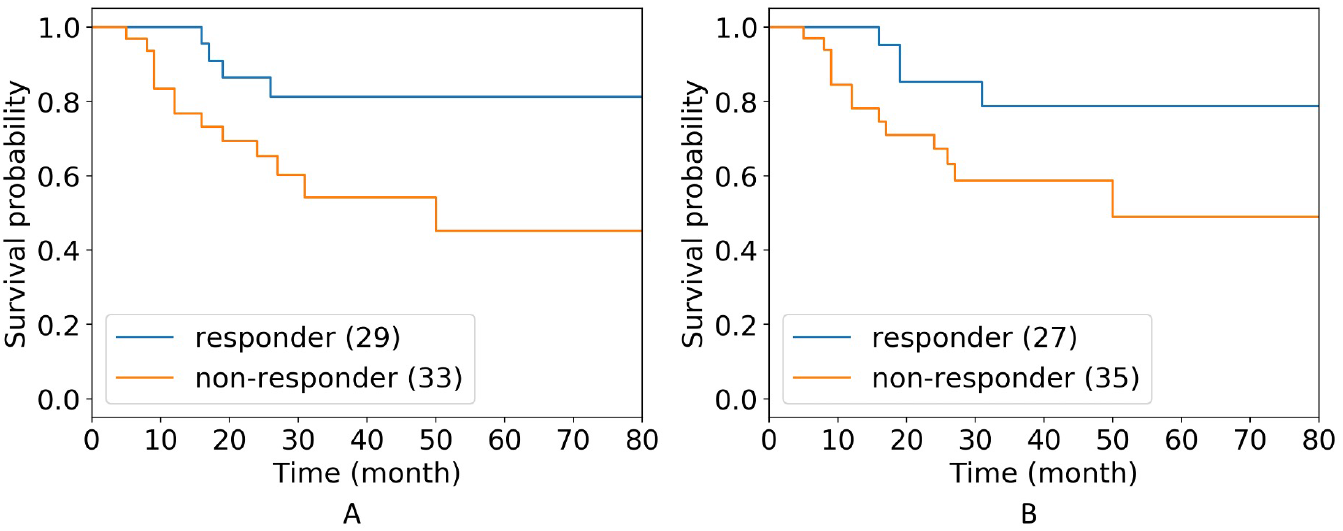
Kaplan-Meier curves of *responder* and *non-responder* group of lung cancer patients that took Cisplatin, Pemetrexed, Paclitaxel, and/or Vinorelbine for adjuvant therapy which these four drugs treated as existing drugs (A) or as new drugs (B).

## 4. Discussion

Accurate prediction of drug sensitivity is crucial for the success of precision oncology. We presented a model called DeepGRMF, which demonstrated enhanced capability for predicting drug sensitivities in comparison to the previous state-of-art algorithm. Furthermore, our model can predict drug responses *de novo* for previously unseen drugs, which enables one to repurpose existing FDA-approved drugs for treating cancer as well as potentially discover novel anticancer chemicals. This capability is achieved by innovative integration of four machine learning technologies: 1) The IDE representation of drugs that incorporates the information of drug chemical structures and MOAs. 2) Matrix-factorization-based collaborative filtering, which captures characteristic interactions between a set of similar cell lines and a set of similar drugs. 3) Graph-based regularization that encodes the similarity of cells and drugs in original input space. As shown in Table A2, using both similarities can improve AUROC and AUPR compared with using one similarity or not using any similarity to constrain solutions. 4) Neural networks that accurately map a new input (a new cell or a drug) to factor space. We also tried other methods to learn the mapping function, such as random forest and elastic net. As shown in Table A3, since the neural network had higher performance, we selected it to learn the mapping functions.

We have shown the generalizability and robustness of our model in transferring the prediction models trained with GDSC data to make predictions on cell lines from another large-scale cell-line-based pharmacogenomic study, CCLE, and, more excitingly, predictions on real-world patients. It is foreseeable that future precision oncology may involve designing a personalized regimen consisting of multiple effective drugs for each patient. Developing prediction systems transferring knowledge from cell lines to real-world clinical practice offers promise for greatly accelerating this process.

The DeepGRMF model can be improved in several aspects. Currently, the DeepGRMF model only utilizes gene expression profiling data, and we anticipate that further integrating genomic alterations (mutations and copy number alterations) and epigenetic information will likely further improve the performance of the model. We noted that the performance of new drug sensitivity prediction is not as good as new cell line prediction, which may be due to limited expressiveness of our VAE-based representation of chemical structures. Recently, it has been shown that a family of graph neural networks can better provide a representation of chemical structures,^26^ which can be explored in the future. Finally, the current representations of cellular states of cancer cells are derived using “black-box” neural networks, and interpretable deep learning models can be explored not only to achieve interpretability of our model but also to enhance its performance.

## Acknowledgment

This work has been partially supported by the grant R01LM012011 from NIH and the AWS ML Research Awards program. Y.T. was partially supported by CMLH Fellowship in Digital Health. R.S. was supported in part by Pennsylvania Dept. of Health award FP00003273 and by the National Human Genome Research Institute of the National Institutes of Health under award number R01HG010589. The content is solely the responsibility of the authors and does not necessarily represent the official views of the above funding agencies. The Pennsylvania Department of Health specifically disclaims responsibility for any analyses, interpretations or conclusions. The results published here are in whole or part based upon data generated by TCGA managed by the NCI and NHGRI. Information about TCGA can be found at http://cancergenome.nih.gov. Preprint of an article published in Pacific Symposium on Biocomputing © 2021 World Scientific Publishing Co., Singapore, http://psb.stanford.edu/.

## 5. Supplementary Information

### 5.1. Implementation

We implemented the DeepGRMF models in Python 3.7.4 with PyTorch. We used grid search for hyperparameter tuning. For training graph regularized matrix factorization, we set the learning rate to 0.001 and mini-batch size as 5096. We set the latent dimension of cell line and drug factor matrix to be 64. Both *λ*_*c*_ and *λ*_*d*_ were 0.2. We considered top 10 and 5 nearest neighbors for cell line and drug, respectively. An early stop was applied after 30 consecutive epochs without loss reduction. For training two neural networks, we used different sets of hyperparameters and model structures due to the difference in feature dimensions of cell line and drug. For the neural network I (*f*_*θ*_), the batch size was 25 with a learning rate 0.00004, and the hidden layers were [2048, 512, 128]. For the neural network II (*f*_*ϕ*_), the batch size was 10, the learning rate was 0.0002 and hidden layers were [192, 128]. We set the dropout rate at 0.2 for both neural networks for each layer except the output layer to prevent overfitting. We also applied early stop for training these two neural networks. For both matrix factorization and neural networks, we used Adaptive Moment Estimation as our gradient descent optimization algorithm. We also reduced the learning rate 10 times as loss stopped decreasing for 10 consecutive training epochs.

### 5.2. Comparison between using gene expression and cell line factor as the features of cell line

Learning more informative features of cell lines is extremely significant in drug sensitivity prediction, since it could improve the accuracy of our predictions. We can provide a better representation of cell lines by using neural network I (*f*_*θ*_) trained in our model. DeepGRMF is a separate model, at step II, we trained a mapping function for cell lines that can map the gene expression data to the cell line factor via neural network I (*f*_*θ*_). To predict drug sensitivity of a new cell line, most methods directly utilize the gene expression profiling of the cell lines as input. We can alternatively provide a cell line factor by applying the trained neural network I (*f*_*θ*_) to the gene expression profiling. We used the lasso model to compare these two representations as the features of cell lines. The result is shown in Table A1. The performance of using the cell line factor as the feature is significantly improved for both AUROC and AUPR.

---

**Table A1:** Performance of using original expression data and cell line factor as features of cell lines to predict drug sensitivity of new cell lines.

| Dataset | Features | Per Cell Line |  | Per Drug |  | Micro |  |
| --- | --- | --- | --- | --- | --- | --- | --- |
|  |  | AUROC | AUPR | AUROC | AUPR | AUROC | AUPR |
| GDSC | Gene Expression | 79.1 | 53.8 | 67.1 | 38.2 | 79.3 | 55.4 |
|  | Cell Line Factor | **82.6** | **59.5** | **70.8** | **41.5** | **82.5** | **61.4** |
| CCLE | Gene Expression | 79.2 | 67.5 | 66.2 | 38.2 | 74.1 | 50.5 |
|  | Cell Line Factor | **82.0** | **70.4** | **67.5** | **40.7** | **75.6** | **52.7** |

**Fig. A1:**
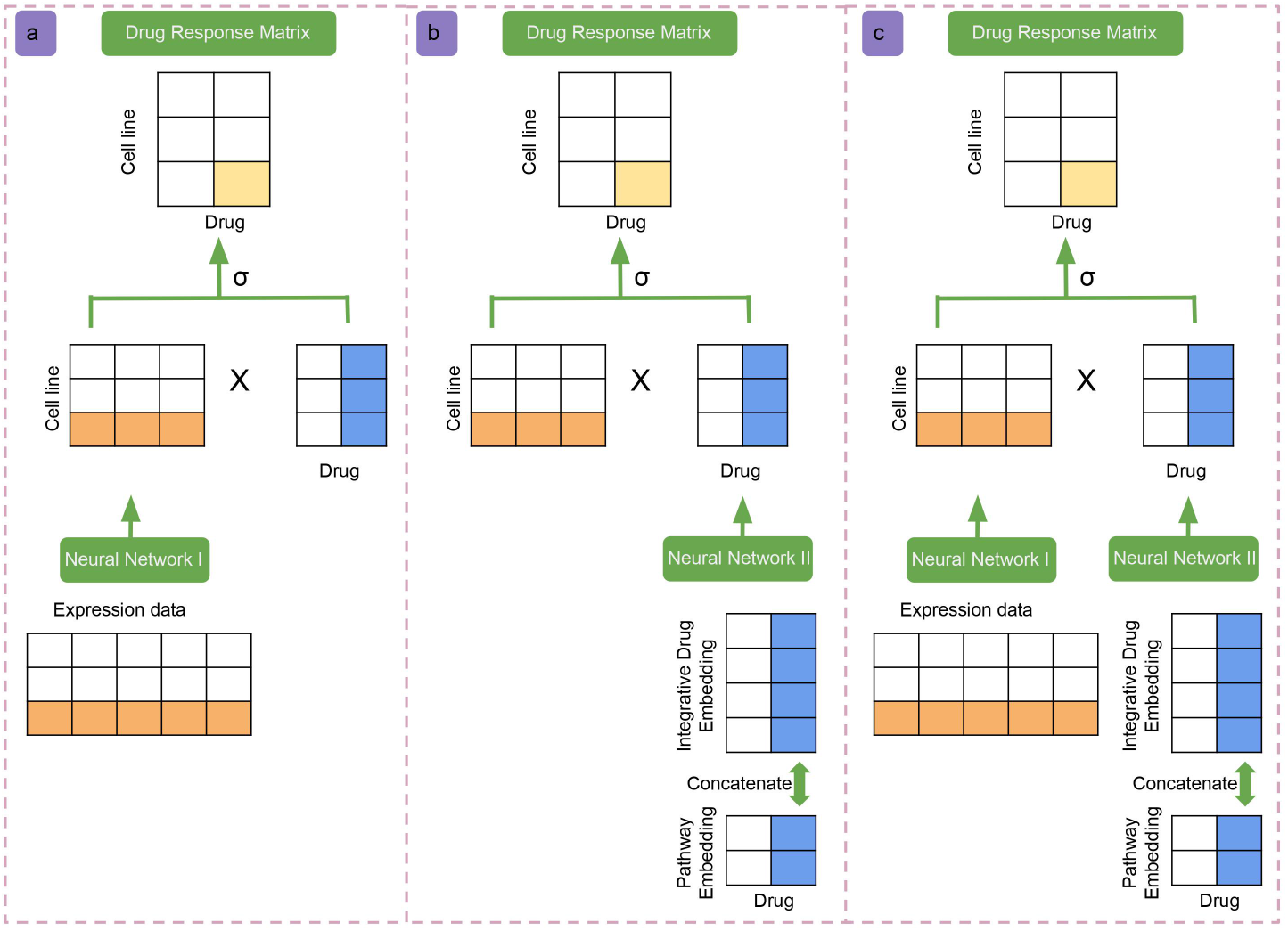
Diagram of DeepGRMF model in three different tasks at the time of testing. Task a is to predict drug response of new cell lines to existing drugs; Task b is to predict drug response of existing cell lines to new drugs; Task c is to predict drug response of new cell lines to new drugs.

**Table A2:**
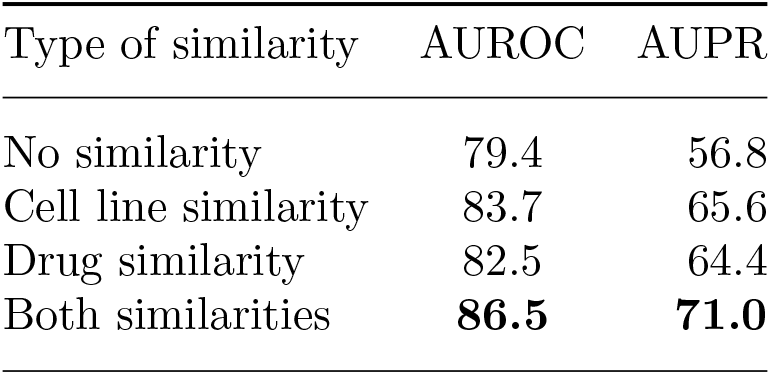
Performance of comparison of using different similarities to regulate the matrix factorization. We calculated the mean AUROC and AUPR in the graph regularized matrix factorization step.

| Type of similarity | AUROC | AUPR |
| --- | --- | --- |
| No similarity | 79.4 | 56.8 |
| Cell line similarity | 83.7 | 65.6 |
| Drug similarity | 82.5 | 64.4 |
| Both similarities | **86.5** | **71.0** |

**Fig. A2:**
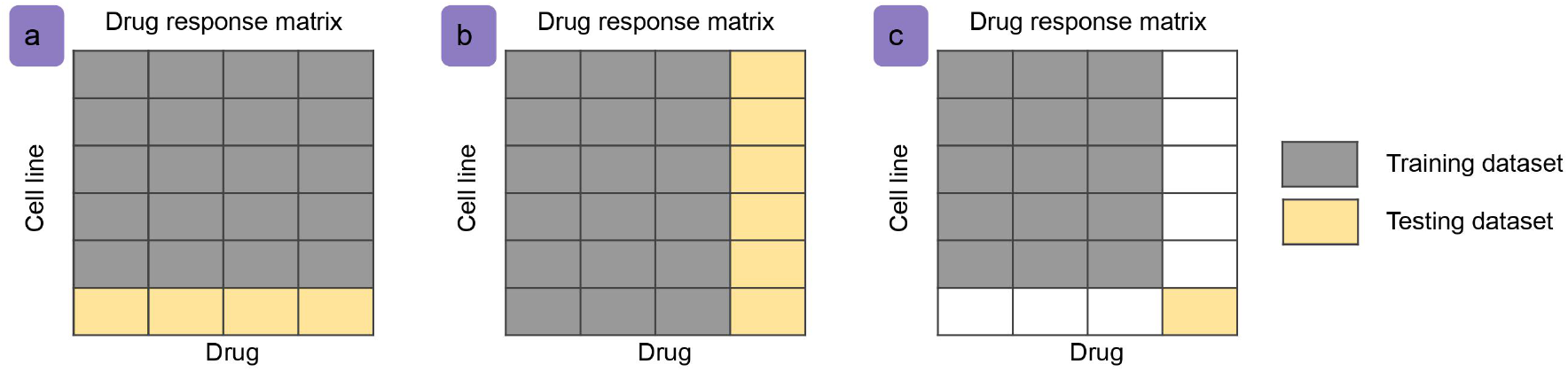
Train-test split strategies in three different tasks. Task a is to predict drug sensitivity of new cell lines to existing drugs; Task b is to predict drug sensitivity of existing cell lines to new drugs; Task c is to predict drug sensitivity of new cell lines to existing drugs.

**Table A3:** Performance of comparison of using different models to map gene expression data to cell line factor space to predict drug sensitivity of new cell lines to existing drugs.

| Dataset | Model | Per Cell Line |  | Per Drug |  | Micro |  |
| --- | --- | --- | --- | --- | --- | --- | --- |
|  |  | AUROC | AUPR | AUROC | AUPR | AUROC | AUPR |
| GDSC | Random Forest | 82.5 | 58.7 | 68.5 | 39.5 | 82.2 | 60.1 |
|  | Elastic Net | 83.0 | 60.0 | 70.1 | 40.7 | 82.8 | 61.4 |
|  | Neural Network | **83.2** | **60.1** | **70.9** | **41.8** | **83.1** | **62.0** |

## Footnotes

* The code and Supplementary Information are available at: https://github.com/renshuangxia/DeepGRMF

## References

1. V. Gambardella, N. Tarazona, J. M. Cejalvo, P. Lombardi, M. Huerta, S. Roselló, T. Fleitas, D. Roda and A. Cervantes, Personalized medicine: recent progress in cancer therapy, Cancers 12, p. 1009 (2020).

2. J. Marquart, E. Y. Chen and V. Prasad, Estimation of the percentage of us patients with cancer who benefit from genome-driven oncology, JAMA oncology 4, 1093 (2018).

3. M. R. Lackner, T. R. Wilson and J. Settleman, Mechanisms of acquired resistance to targeted cancer therapies, Future oncology 8, 999 (2012).

4. Y. Tao, H. Lei, X. Fu, A. V. Lee, J. Ma and R. Schwartz, Robust and accurate deconvolution of tumor populations uncovers evolutionary mechanisms of breast cancer metastasis, Bioinformatics 36, i407 (07 2020).

5. W. Yang, J. Soares, P. Greninger, E. J. Edelman, H. Lightfoot, S. Forbes, N. Bindal, D. Beare, J. A. Smith, I. R. Thompson et al., Genomics of drug sensitivity in cancer (gdsc): a resource for therapeutic biomarker discovery in cancer cells, Nucleic acids research 41, D955 (2012).

6. J. Barretina, G. Caponigro, N. Stransky, K. Venkatesan, A. A. Margolin, S. Kim, C. J. Wilson, J. Lehár, G. V. Kryukov, D. Sonkin et al., The cancer cell line encyclopedia enables predictive modelling of anticancer drug sensitivity, Nature 483, 603 (2012).

7. K. Tomczak, P. Czerwińska and M. Wiznerowicz, The cancer genome atlas (tcga): an immeasurable source of knowledge, Contemporary oncology 19, p. A68 (2015).

8. A. Subramanian, R. Narayan, S. M. Corsello, D. D. Peck, T. E. Natoli, X. Lu, J. Gould, J. F. Davis, A. A. Tubelli, J. K. Asiedu et al., A next generation connectivity map: L1000 platform and the first 1,000,000 profiles, Cell 171, 1437 (2017).

9. A. B. Keenan, S. L. Jenkins, K. M. Jagodnik, S. Koplev, E. He, D. Torre, Z. Wang, A. B. Dohlman, M. C. Silverstein, A. Lachmann et al., The library of integrated network-based cellular signatures nih program: system-level cataloging of human cells response to perturbations, Cell systems 6, 13 (2018).

10. X. Su and T. M. Khoshgoftaar, A survey of collaborative filtering techniques, Advances in artificial intelligence 2009 (2009).

11. M. Q. Ding, L. Chen, G. F. Cooper, J. D. Young and X. Lu, Precision oncology beyond targeted therapy: combining omics data with machine learning matches the majority of cancer cells to effective therapeutics, Molecular cancer research 16, 269 (2018).

12. Y.-C. Chiu, H.-I. H. Chen, T. Zhang, S. Zhang, A. Gorthi, L.-J. Wang, Y. Huang and Y. Chen, Predicting drug response of tumors from integrated genomic profiles by deep neural networks, BMC medical genomics 12, 143 (2019).

13. Y. Tao, S. Ren, M. Q. Ding, R. Schwartz and X. Lu, Predicting drug sensitivity of cancer cell lines via collaborative filtering with contextual attention, Proceedings of Machine Learning Research Vol. 126 (PMLR, Virtual, 07–08 Aug 2020).

14. J. Cadow, J. Born, M. Manica, A. Oskooei and M. Rodríguez Martínez, PaccMann: a web service for interpretable anticancer compound sensitivity prediction, Nucleic acids research 48, W502 (2020).

15. M. Li, Y. Wang, R. Zheng, X. Shi, Y. Li, F.-X. Wu and J. Wang, DeepDSC: A deep learning method to predict drug sensitivity of cancer cell lines, IEEE/ACM transactions on computational biology and bioinformatics 18, 575 (2021).

16. M. P. Menden, F. Iorio, M. Garnett, U. McDermott, C. H. Benes, P. J. Ballester and J. Saez-Rodriguez, Machine learning prediction of cancer cell sensitivity to drugs based on genomic and chemical properties, PLoS one 8, p. e61318 (2013).

17. Y. Xue, M. Q. Ding and X. Lu, Learning to encode cellular responses to systematic perturbations with deep generative models, NPJ systems biology and applications 6, 1 (2020).

18. J. Zhu, J. Wang, X. Wang, M. Gao, B. Guo, M. Gao, J. Liu, Y. Yu, L. Wang, W. Kong, Y. An, Z. Liu, X. Sun, Z. Huang, H. Zhou, N. Zhang, R. Zheng and Z. Xie, Prediction of drug efficacy from transcriptional profiles with deep learning, Nature biotechnology (2021).

19. K. Yu, S. Visweswaran and K. Batmanghelich, Semi-supervised hierarchical drug embedding in hyperbolic space, Journal of chemical information and modeling 60, 5647 (2020).

20. D. S. Wishart, C. Knox, A. C. Guo, S. Shrivastava, M. Hassanali, P. Stothard, Z. Chang and J. Woolsey, DrugBank: a comprehensive resource for in silico drug discovery and exploration, Nucleic acids research 34, D668 (2006).

21. N.-N. Guan, Y. Zhao, C.-C. Wang, J.-Q. Li, X. Chen and X. Piao, Anticancer drug response prediction in cell lines using weighted graph regularized matrix factorization, Molecular therapy-nucleic acids 17, 164 (2019).

22. D. Weininger, SMILES, a chemical language and information system. 1. introduction to methodology and encoding rules, Journal of chemical information and computer sciences 28, 31 (1988).

23. J. J. Irwin, T. Sterling, M. M. Mysinger, E. S. Bolstad and R. G. Coleman, ZINC: a free tool to discover chemistry for biology, Journal of chemical information and modeling 52, 1757 (2012).

24. D. P. Kingma and M. Welling, Auto-encoding variational bayes, in 2nd International Conference on Learning Representations, (Banff, AB, Canada, 2014).

25. J. Xu, V. Jagadeesh and B. Manjunath, Multi-label learning with fused multimodal bi-relational graph, IEEE transactions on multimedia 16, 403 (2014).

26. M. Sakai, K. Nagayasu, N. Shibui, C. Andoh, K. Takayama, H. Shirakawa and S. Kaneko, Prediction of pharmacological activities from chemical structures with graph convolutional neural networks, Scientific reports 11, 1 (2021).

